# AlphaBind, a Domain-Specific Model to Predict and Optimize Antibody-Antigen Binding Affinity

**DOI:** 10.1101/2024.11.11.622872

**Authors:** Aditya A. Agarwal, James Harrang, David Noble, Kerry L. McGowan, Adrian W. Lange, Emily Engelhart, Miranda C. Lahman, Jeffrey Adamo, Xin Yu, Oliver Serang, Kyle J. Minch, Kimberly Y. Wellman, David A. Younger, Randolph M. Lopez, Ryan O. Emerson

## Abstract

Antibodies are versatile therapeutic molecules that utilize combinatorial sequence diversity to cover a vast fitness landscape. However, designing optimal antibody sequences remains a major challenge. Recent advances in deep learning provide opportunities to address this challenge by learning sequence-function relationships to accurately predict fitness landscapes. These models enable efficient *in silico* prescreening and optimization of antibody candidates. By focusing experimental efforts on the most promising candidates guided by deep learning predictions, antibodies with optimal properties can be designed more quickly and effectively.

Here we present AlphaBind, a domain-specific model that utilizes protein language model embeddings and pre-training on millions of quantitative laboratory measurements of antibody-antigen binding strength to achieve state-of-the-art performance for guided affinity optimization of parental antibodies. We demonstrate that an AlphaBind-powered antibody optimization pipeline can deliver candidates with substantially improved binding affinity across four parental antibodies (some of which were already affinity-matured) and using two different types of training data. Resulting candidates, ranging up to 11 mutations from parental sequence, yield a sequence diversity that allows for optimization of other biophysical characteristics, all while using only a single round of data generation for each parental antibody. AlphaBind weights and code are publicly available at: https://github.com/A-Alpha-Bio/alphabind.

## Introduction

Therapeutic antibodies play an increasingly pivotal role in the biotherapeutics market, and the ability for antibody variable fragments to participate in high-affinity and highly specific protein-protein interactions makes them ideal for therapeutic modalities including antibody-drug conjugates, bispecific antibodies, CAR-T therapies, etc. Their development has historically relied on labor-intensive experimental methods including directed evolution via display techniques or via animal immunization, followed by manual optimization through iterative assessment of a small number of human-designed variants^1,2^.

For the development of such molecules, it is essential to control and optimize the binding affinity of an antibody to its target, sometimes to multiple targets, while also ensuring maintenance of a valid antibody-like structure and favorable biochemical properties like solubility, aggregation and potential immunogenicity^3-6^. However, the plausible search space is both enormous (e.g., each amino acid change from a parental antibody of length 100 increases the search space by a factor of approximately 2,000) and sparse (most sequences derived from any given parental antibody will be poor binders, poorly behaved as recombinant protein, or both^7^). The challenge is increased when a candidate antibody must bind more than one target (e.g., cross-reactivity across a phylogeny of target sequences in an infectious disease context), or when properties other than affinity make orthogonal demands on primary sequence (e.g. multi-objective optimization for affinity plus solubility, thermostability, etc.).

Recently, deep learning methods have greatly improved the ability to effectively sample valid antibody sequences, including evolution- and structure-informed approaches, and both unguided methods, and guided methods incorporating additional *in vitro* data^8-11^. Among these approaches, effective model scaling has been demonstrated according to model complexity as well as the amount and quality of training data^12-14^. Unguided generative sampling of valid sequences near a parental sequence can produce proposal sets, including many that can be expressed as functional proteins and may have similar or improved function compared to the parental sequence^15-17^. Given any such technique for sampling valid sequences, design of a novel therapeutic sequence can be further accelerated via prioritizing proposals for *in vitro* validation based on an appropriate fitness landscape—assuming a suitable prediction of fitness is available. Taking this guided approach, several studies have demonstrated the effectiveness of deep learning approaches incorporating large, high-quality datasets to train highly effective affinity prediction models used to prioritize candidate sequences for *in vitro* validation^18-20^.

The AlphaSeq assay^21^, a yeast display system that quantitatively measures large numbers of protein-protein interaction affinities in parallel, has been leveraged to generate such datasets characterizing antibody affinity using large libraries of antibody and antigen sequence variants, including a large dataset of largely scFv and VHH antibodies vs. a SARS-CoV-2 RBD mutation library^22^; a dataset of scFv-format antibodies vs. a SARS-CoV-2-derived peptide^23^; and several large panels comprising variants of VHH72, a camelid antibody targeting SARS-CoV-1 RBD with some SARS-CoV-2 RBD cross-reactivity^18^. Multiple studies using these datasets have previously demonstrated that AlphaSeq data are highly effective for training machine learning models for guided antibody sequence optimization^18,19^.

Here, we combine high-throughput antibody-antigen affinity datasets which explore the local fitness landscape around a parental antibody with AlphaBind, a domain-specific model pre-trained on approximately 7.5 million quantitative affinity measurements from unrelated antibody-antigen systems, to perform guided antibody optimization on three diverse parental antibodies using AlphaSeq datasets to fine-tune our pretrained model. We demonstrate that (1) given adequate local training data, sequence optimization using AlphaBind quickly generates thousands of high-affinity antibody derivatives, (2) both protein language model embeddings and pre-training on unrelated antibody-antigen binding data contribute significantly to model performance, and (3) the AlphaBind model retains good performance even on local data generated via a different assay than its own training data, using previously published high throughout mammalian display data exploring CDRH3 mutations of trastuzumab to conduct a fourth optimization campaign. Furthermore, we show that a fine-tuned AlphaBind model can guide sequence optimization for predicted developability and immunogenicity while maintaining affinity, resulting in an optimized variant of our anti-TIGIT scFv with no predicted sequence liabilities and extensive sequence reversion to human germline with no loss in affinity. Finally, we present the necessary background, model architecture and code, including a pre-trained model checkpoint, to enable broad application of the AlphaBind model for affinity optimization of therapeutic antibody candidates.

## Results

### Antibody Optimization Campaigns

Three different antibody-antigen systems were chosen for benchmarking of AlphaSeq-based antibody optimization with AlphaBind: AAB-PP489, a previously-undescribed humanoid scFv targeting human TIGIT; Pembrolizumab-scFv, an scFv-formatted humanized mouse antibody targeting human PD-1^24^; and VHH72, a camelid single-chain antibody targeting the RBD of SARS-CoV1 with cross-reactivity to SARS-CoV2^25^. Figure 1 provides an overview of the AlphaBind fine-tuning and optimization process, and additional details for each system are given in Table S1.

**Figure 1.**
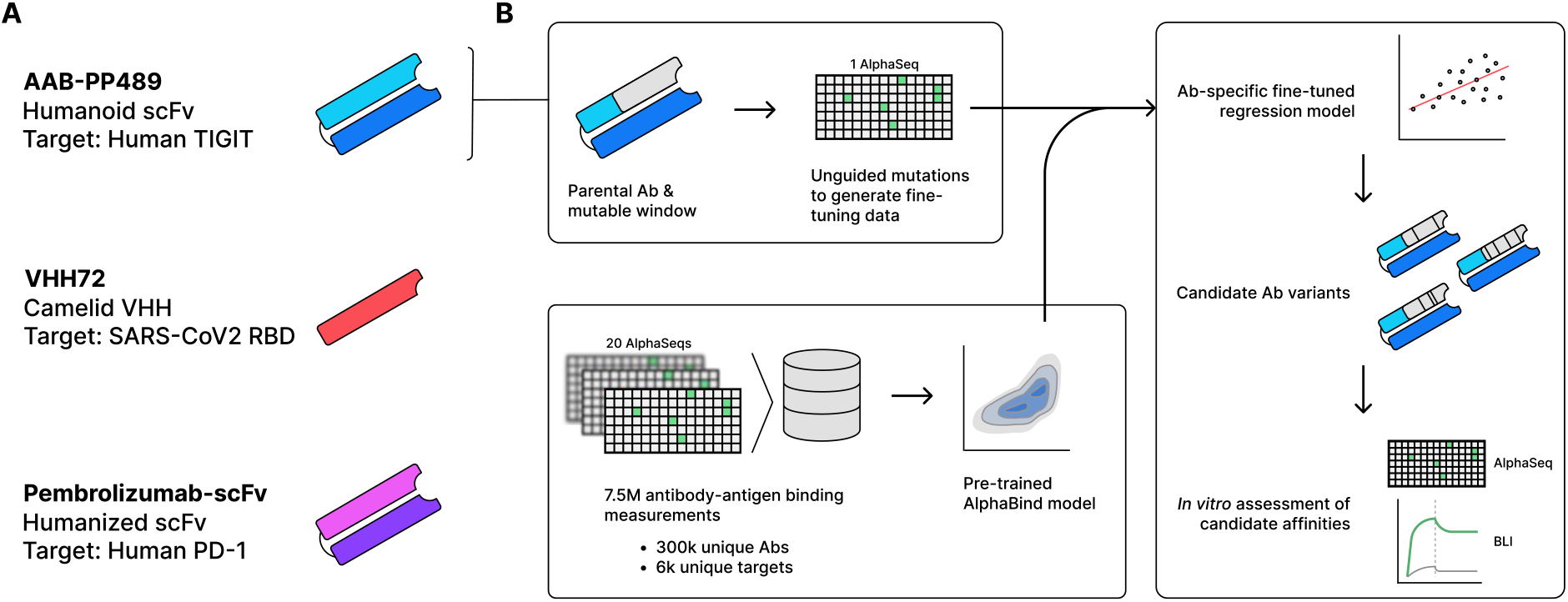
Experimental Overview. **A**: Summary of the three parental antibodies used to benchmark AlphaBind, including a humanoid scFv binding human TIGIT (AAB-PP489), a camelid single-domain heavy chain binding SARS-CoV2 RBD (VHH72), and a humanized mouse scFv binding human PD-1 (Pembrolizumab-scFv). **B:** For each parental antibody, a contiguous 300nt region (including 5’ and 3’ primer sites) was identified, defining the mutable window; AlphaSeq was used to generate a training dataset of approximately 30,000 variants within the mutable window. The AlphaBind regressor, pre-trained on unrelated antibody-antigen affinity data, was fine-tuned with each training dataset to generate an antibody-specific sequence-to-affinity regressor, and then used to generate candidate sequences optimized for predicted binding affinity. Candidate variants of each parental antibody were validated *in vitro* using AlphaSeq for large-scale validation and biolayer interferometry for a subset of candidates.

For each parental antibody, a fine-tuning dataset was generated by constructing a library of approximately 30,000 antibody variants. A contiguous region of approximately 83 amino-acid residues was selected for each antibody covering as much of the CDRH regions as possible; limiting the mutable region to 83 amino acids allowed generation of large variant libraries using 300-nt oligonucleotides including fixed flanks for cloning. Within each mutable region, we constructed a yeast display library comprising essentially all single missense mutations (except for cysteine), a limited number of insertions and deletions, many randomly selected double and triple missense mutations, and several replicates of the parental antibody sequence. These libraries were then used to generate AlphaSeq data, providing quantitative affinity values for binding between each library member and its associated target; across targets, most variants had measurable binding to their targets with a minority showing improved affinity, establishing the quantitative data necessary to learn an antibody-specific fitness landscape for each parental antibody. Fine-tuning AlphaSeq datasets are available as part of the Supplemental Materials, and Table S2 provides a summary of datasets generated as part of this work.

### Model Training & Optimization

Figure 2 summarizes the architecture and pre-training of the AlphaBind regression model and our optimization strategy. Briefly, the AlphaBind model was pre-trained on antibody-antigen AlphaSeq affinity data from unrelated systems. Twenty distinct AlphaSeq datasets were used for pre-training, comprising approximately 7.5 million rows of antibody-antigen affinity data, including roughly 300,000 distinct antibody constructs and 6,000 distinct target constructs. The pre-training data included variant libraries, comprising tens of thousands of variants from various parental antibodies, as well as diverse libraries such as libraries cloned from phage panning experiments— thus including tens of thousands of distinct parental antibodies. Training data were screened to prevent inclusion of any data related to the parental antibodies or targets being optimized herein. As a result, only Pembrolizumab-scFv used the full pre-trained model for fine-tuning—AAB-PP489 and VHH72 utilized a smaller pre-trained model (see Materials and Methods).

**Figure 2.**
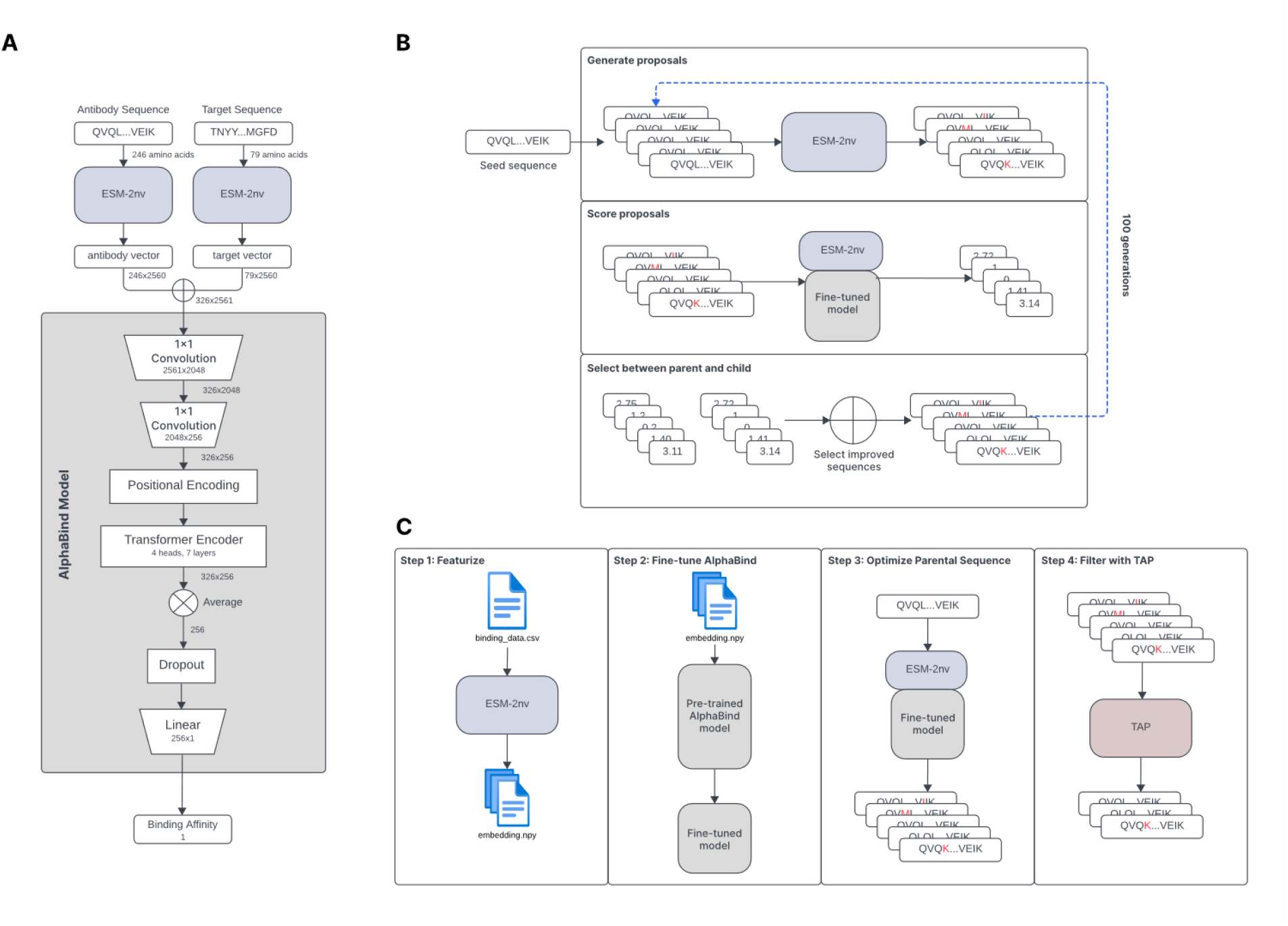
AlphaBind Architecture. **A:** Antibody and target are encoded using ESM-2nv, then those tensors are concatenated and input into the AlphaBind model, including a transformer encoder with 4 heads and 7 layers and a total of 15M parameters. **B:** During optimization, ESM-2nv masking is used to perturb seed sequences, which are scored according to the fine-tuned regression model and accepted if they have improved predicted affinity, then the process is iterated. **C:** Overall campaign architecture: training data are featurized, the AlphaBind regressor is fine-tuned, optimization is performed to generate a pool of candidates, and TAP^5,6^ is used to filter the resulting candidates to ensure no major predicted developability issues.

After pre-training, AlphaSeq datasets for each parental antibody were used to fine-tune the AlphaBind model. Each fine-tuned regression model was used to generate approximately 6 million sequence proposals, from which approximately 7,500 candidate sequences were selected for *in vitro* validation in AlphaSeq. To explore AlphaBind’s ability to extrapolate beyond its training data, candidates were selected according to a uniform distribution for 2 through 11 mutations with respect to the parental antibody. Among these 7,500 candidates, the top 5 candidates with the strongest predicted affinity were selected for expression and validation using BLI, without regard to number of mutations. See Materials and Methods for detailed information on model training, optimization, and sequence selection.

An analysis of sequence diversity among the 7,500 candidates chosen for AlphaSeq validation for each parental antibody demonstrates a broad range of both positions mutated and variety of mutations selected, as shown in Figure 3 (CDR regions annotated using ANARCI, according to Chothia nomenclature)^26,27^. For AAB-PP489, AlphaBind preferred to mutate CDRH3 (as might be expected for a phage panning hit not otherwise affinity-matured to a local optimum), while also making many mutations in FR3. For VHH72, diversity was highest in CDRH1 and CDRH2, also with many mutations in FR3. For Pembrolizumab-scFv, only a single position in CDRH3 had high sequence diversity among optimized candidates, again with many mutations made in FR3. This analysis demonstrates that AlphaBind candidates cover a wide sequence space, providing substantial freedom for downstream selection of additional properties such as predicted biophysical properties.

**Figure 3.**
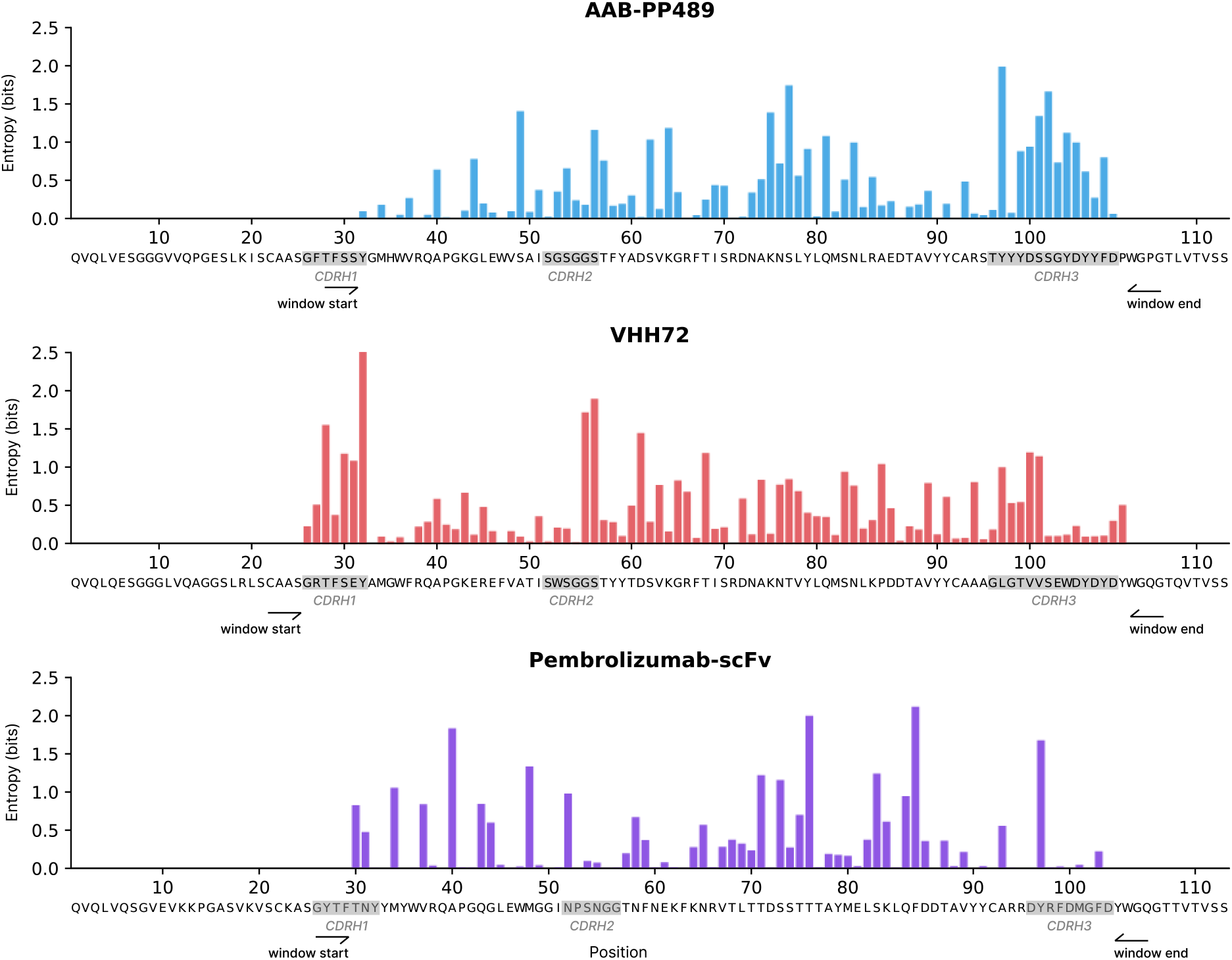
Sequence Diversity of AlphaSeq Optimized Candidates. For each parental antibody, we annotate CDRH regions and plot the primary sequence diversity (Shannon entropy, measured in bits) at each position within the designated mutational windows. While there is substantial variation in the mutability of each position, AlphaBind proposes a broad variety of mutations throughout the CDR and framework regions, with many different positions mutated and many different mutations observed at key positions.

### *In Vitro* Validation of AlphaBind Candidates

AlphaSeq validation results for each parental antibody are summarized in Figure 4A; across all parental antibodies, AlphaBind fine-tuning and optimization was able to generate thousands of candidates with binding affinity stronger than the parental. Up to 8 mutations away from the parental for VHH72, and up to 11 mutations away from parental (the maximum tested) for Pembrolizumab-scFv and AAB-PP489, the median binding affinity for an AlphaBind-proposed candidate was better than its parent. Across campaigns, candidates were specific for their intended targets: among candidates with ≤ 1 µM AlphaSeq affinity to their target, 80% of VHH72 candidates, 92% of Pembrolizumab-scFv candidates, and 97% of AAB-PP489 candidates demonstrated specific binding defined as on-target affinity ≥ 100x higher than average off-target affinity measured to irrelevant control proteins in AlphaSeq.

**Figure 4.**
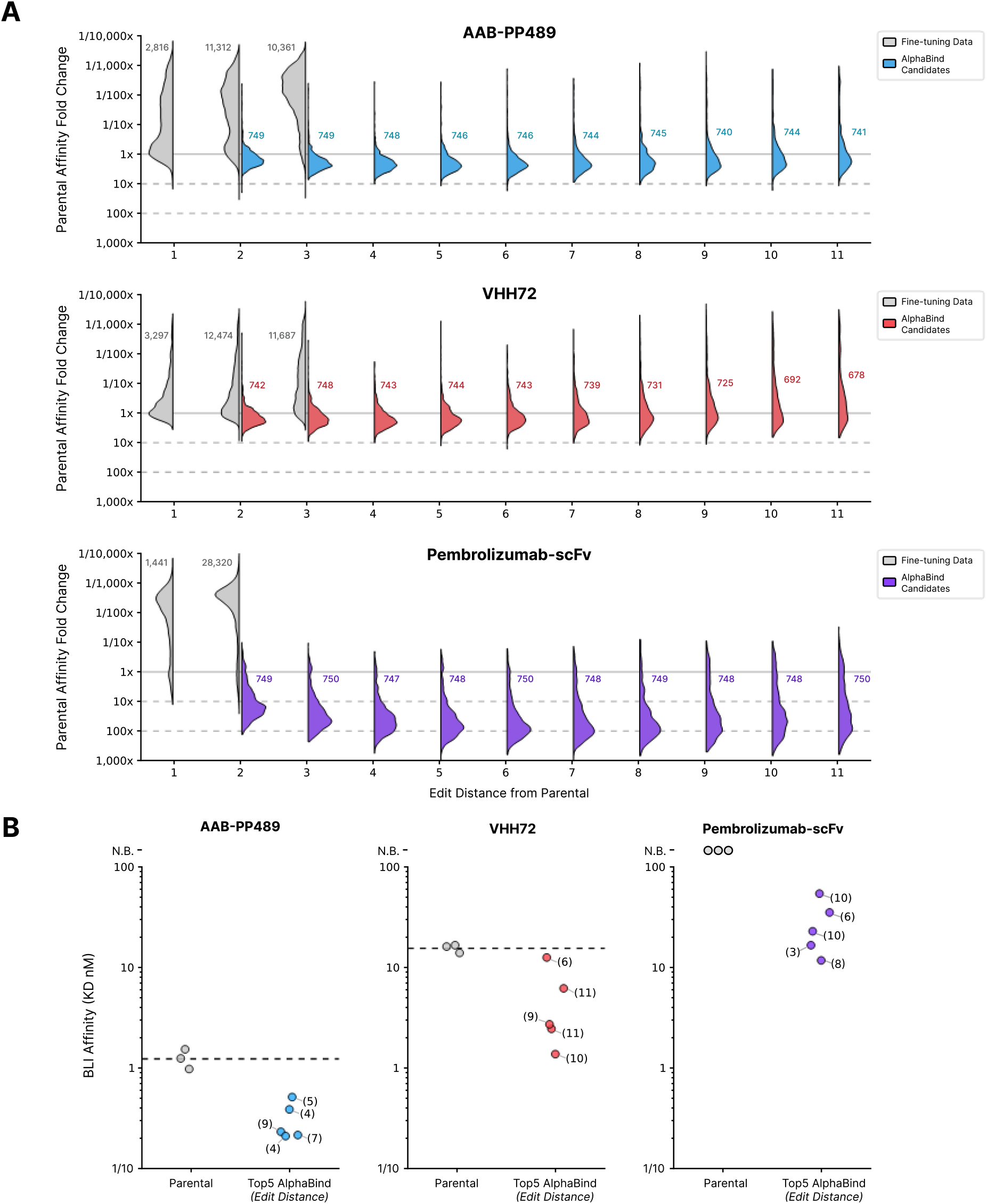
AlphaSeq Validation Results. **A:** Fine-tuned AlphaBind models were trained for each of three parental antibodies and used to generate 750 candidates at each edit distance from 2 through 11 mutations relative to parental. AlphaSeq was then used to prospectively assess the affinity of optimized candidates. For each parental antibody, we report the distribution of affinities for optimized candidates at each edit distance as affinity relative to the mean affinity among 50 parental antibody replicate measurements. Numerical annotations represent the number of candidates tested at each edit distance. AlphaBind-derived candidates have a median affinity better than the parental molecule up to 8 edits away from the parent sequence for VHH72 and up to 11 edits from the parent sequence for Pembrolizumab-scFv and AAB-PP489. **B:** BLI validation results for the top 5 AlphaBind candidates for each parental antibody. Mean parental affinities are displayed as dashed lines. Across the three systems, 15/15 (100%) top candidates were successfully expressed, and of those 10/10 (100%) had better affinity than the parental antibody. Affinity relative to parental antibody could not be assessed for Pembrolizumab-scFv since the parental antibody was not successfully expressed and thus no BLI measurement was available. No candidates had any measurable binding to an irrelevant negative control target. Single-point BLI values for certain candidates were confirmed via either multi-point BLI or KinExA for candidates with off-rates at or near the BLI limit of detection, confirming a 74x improvement in affinity for the best AAB-PP489 candidate and a 14x improved affinity for the best VHH72 candidate.

Figure 4B summarizes the results of the top 15 optimized candidates selected for BLI validation (5 per parental antibody); 15 out of 15 candidates (100%) were successfully expressed as scFv-FC, and 10 out of 10 (100%) had superior binding affinity than their parental antibodies. The parental Pembrolizumab-scFv could not be expressed in scFv-FC format; all 5 top variants did express but affinity could therefore not be compared to parental. No candidates had any binding to an irrelevant negative control target by BLI. Affinity for the top VHH72 candidates was confirmed by multi-point BLI. Several AAB-PP489 candidates had binding affinities outside the accurate range of BLI and failed to generate a reliable off-rate (i.e., reported as < 1.0E-4/s). Affinity for the top performing AAB-PP489 variant and matched parental was confirmed via kinetic exclusion assay (KinExA). We confirmed an affinity improvement of 74x for the best AAB-PP489 variant (from 56.5 pM for parental to 766 fM, by KinExA) and 14x for the best VHH72 variant (from 16.8 nM to 1.17 nM, by multi-point BLI).

### Affinity-Guided Developability Engineering

To demonstrate the utility of a fine-tuned AlphaBind affinity model for downstream engineering, we used the fine-tuned AAB-PP489 model to design variants with improved developability attributes. We selected AAB-PP3115, an affinity-optimized variant of AAB-PP489, for further engineering due to its strong affinity and observed biophysical properties (see Supplemental Materials for the sequence of AAB-PP3115 and its variants). This engineering campaign is summarized in Figure 5; briefly, AAB-PP3115 contains two potential sequence liabilities, a DS isomerization motif and a DP fragmentation motif in CDRH3, and has nine residues that do not match human germline (IGHV3-23*04 and IGHJ5*02) within its heavy chain mutable window. Our goal was to ablate those sequence liabilities (improving predicted developability) and revert as many residues as possible to germline (improving predicted immunogenicity) while maintaining or improving affinity to human TIGIT. First, we generated (19^4^ = 130,321) candidates with all possible mutations to the four residues involved in the liability motifs, keeping 736 that removed both predicted liabilities and had an AlphaBind-predicted affinity within 0.5 logs of parental AAB-PP3115. Then, for each of those 736 we generated 2^9^ (512) sequences with all combinations of germline-reverted residues, for a total of 376,832 candidates. From these candidates we selected 11 with the most germline reversions, best AlphaBind-predicted affinity, and favorable predicted developability and immunogenicity metrics according to TAP and NaturalAntibody^5,28^, then expressed those candidates for *in vitro* validation alongside parental AAB-PP3115. As a control for the trivial case of full germline reversion, we expressed one candidate with the best predicted liability-reversion mutations and all nine positions reverted to germline, predicted by AlphaBind to have a 337-fold reduced affinity compared to parental. This process led to the identification of AAB-PP3117, with no liability motifs in the mutable window, 4 of 9 residues reverted to human germline, improved expression compared to its parental molecule (334.1 µg/mL compared to 259.1 µg/mL), and improved binding to human TIGIT. We measured an affinity of 310 fM in IgG format by KinExA, a 182-fold improvement over the original AAB-PP489 (56.5 pM scFv-FC by KinExA), and a 3.4-fold improvement over its parent AAB-PP3115 (1.04 pM IgG by KinExA), despite making 7 additional mutations guided only by the fine-tuned AlphaBind regression model. As predicted, the fully-germlined variant was a poor binder, underscoring the need for ML guidance to maintain affinity when undertaking additional sequence engineering. Details of all tested candidates, as well as full BLI and KinExA results, are available in the Supplemental Materials, including BLI and protein analytics in Table S5 and KinExA results in Table S7.

**Figure 5.**
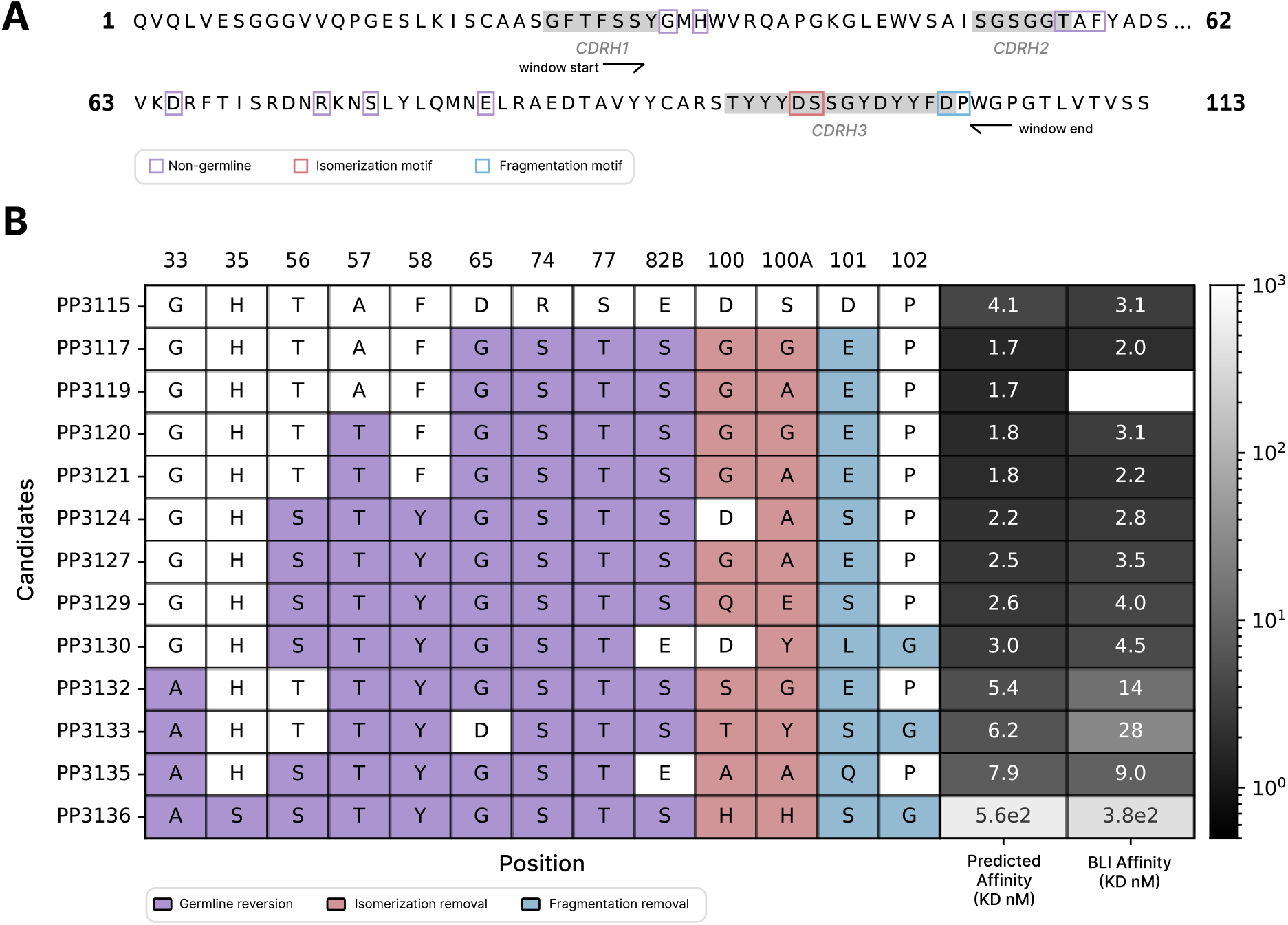
Affinity-Guided Developability Engineering. **A:** AAB-PP3115 is an AlphaBind variant of AAB-PP489 with optimized affinity. Within its mutable window, AAB-PP3115 contains nine non-germline residues along with one isomerization liability motif and one fragmentation liability motif. **B:** The fine-tuned AAB-PP489 AlphaBind regressor was used to assess approximately 500,000 derivatives of AAB-PP3115, in order to ablate an isomerization motif and a fragmentation motif in CDRH3, and revert as much of FR2/FR3 as possible to human germline; 11 candidates were chosen for *in vitro* validation by BLI. 10/11 (91%) candidates expressed, 7/11 (64%) maintained affinity within 2x and 3/11 (27%) had improved affinity for human TIGIT, all while making an additional 7-9 mutations in the successful sequences. Full analytics results including titer, BLI affinity, ASEC and DSF are provided in Table S5.

### Assessment of Ablated AlphaBind Models

To investigate the relative contribution of different AlphaBind architectural components, we performed analogous regressor training and affinity optimization using four ablated versions of the AlphaBind model. Results are summarized in Figure 6, with additional details in Materials and Methods: briefly, we compared the optimized candidate sequences from each ablated model, either in full (all candidates) or among top candidates after *in vitro* validation with AlphaSeq (i.e., just those most likely be chosen for further development). Each ablated model was compared to the full AlphaBind model according to median candidate affinity, with p-values computed via Mann-Whitney U test, assessing the null hypothesis that candidates from the full AlphaBind model had no better affinity than candidates from each ablated model. The simplest transformer-based regression model with one-hot encoded sequences (‘ohe_cold’), while still generating many candidates with improved affinity over parental and extrapolating several mutations farther than its training data, generated candidates much worse than the full AlphaBind model. Two other ablated regressor architectures, one-hot encoded sequences plus pretraining (‘ohe_warm’) and ESM-2nv embedded sequences without pretraining (‘esm_cold’), demonstrated intermediate results with the combination of ESM-2nv-embedded input sequences and pretraining together accounting for AlphaBind performance. An ablated model with an identical regressor but lacking ESM-2nv masking during the optimization step (‘esm_warm’) was nearly equivalent in candidate performance, but substantially increased overall optimization cost (the ablated model had to be optimized for 300 generations, rather than 100 for the full AlphaBind model, to generate sufficient candidates at edit distances of up to 11 mutations). Overall, the full AlphaBind model had significantly improved performance over models which lacked either ESM-2nv embedding or pre-training on related data, with a substantial difference in the quality of the top candidates in particular.

**Figure 6.**
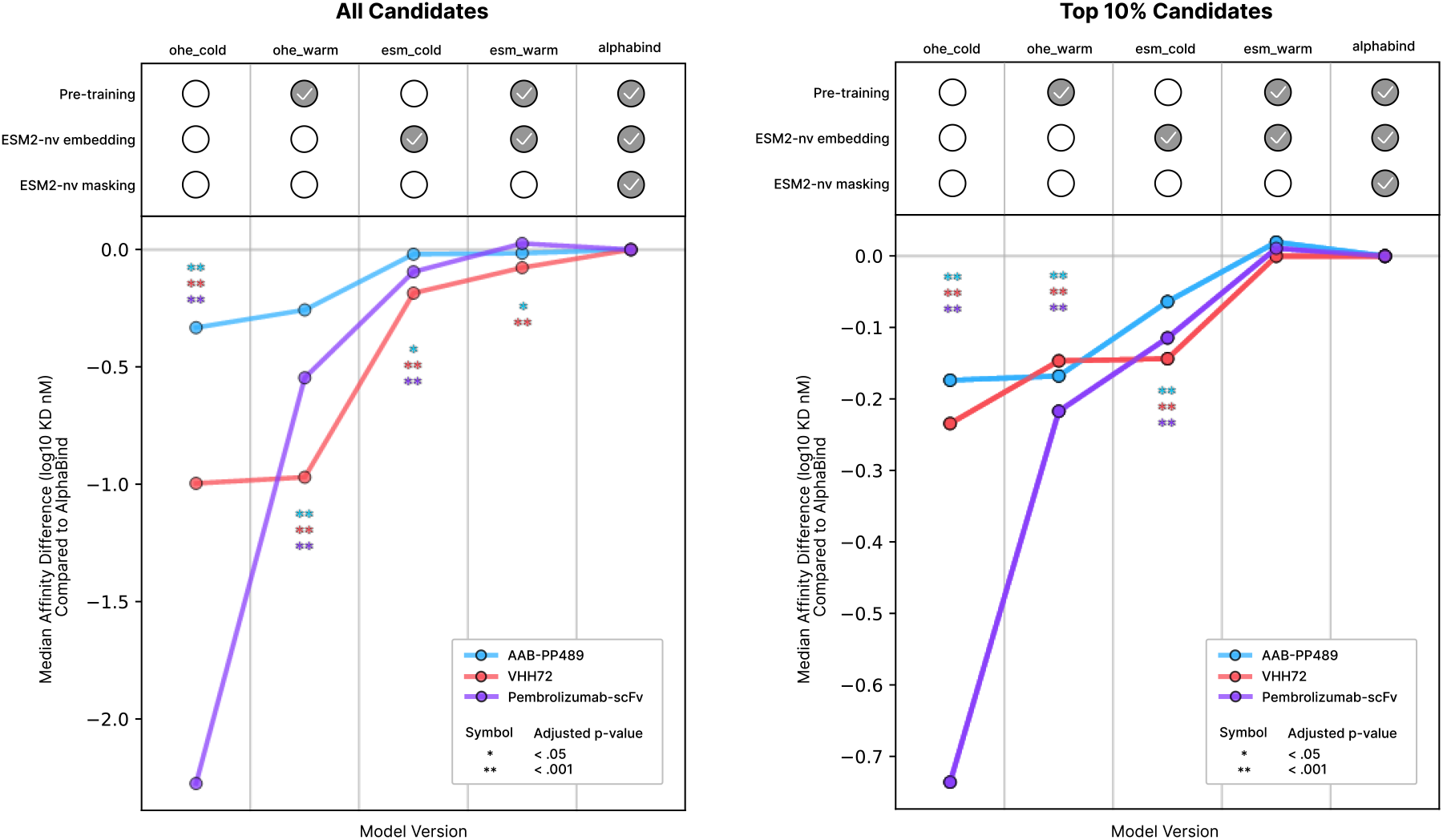
AlphaBind Ablation Analysis. **A:** For each experimental system, all 7,500 candidates were compared for each of 5 models. The difference in median affinity between AlphaBind candidates and candidates from each ablated model is plotted, and significance is calculated according to a Mann-Whitney U test on candidate affinities with Benjamini-Hochberg procedure for control of false discovery. The removal of ESM-2nv embedding or pre-training on unrelated AlphaSeq data both significantly degrade overall model performance, while removal of ESM-2nv masking during optimization is equivalent or marginally superior, though less computationally efficient. **B:** Ablated model performance, according to the same procedure and metrics as above, but focused only on the top 10% of candidates according to AlphaSeq validation, i.e., the candidates most likely to be chosen for additional validation or engineering; again, the effect of ESM-2nv masking during optimization is marginal to slightly negative, but removal of either ESM-2nv embedding or pre-training substantially degrades performance.

### Antibody Optimization with Alternative Fine-Tuning Data

To assess the utility of the AlphaBind pre-trained model with fine-tuning data from sources other than our own AlphaSeq assay, we performed an optimization campaign on Trastuzumab-scFv, a humanized mouse antibody targeting human HER2^29^. Trastuzumab has been a common benchmarking system for the design and optimization of antibody variants using machine learning, with many studies assessing different methods^15,16,20,30^, and variant binding datasets are readily available for fine-tuning. We utilized the affinity-sorted mammalian display dataset of approximately 36,000 CDRH3 variants from Mason, et al.^20^ as fine-tuning data, and performed two different optimization experiments with the resulting fine-tuned AlphaBind model: first, we optimized just the CDRH3 region, concordant with the domain of the training dataset; second, we allowed our optimization routine to edit a larger contiguous window comprising the CDRH2-FR3-CDRH3 region, with edits to the CDRH2 and FR3 regions representing out-of-distribution prediction. As with the other optimization campaigns, we selected 7,500 candidates for AlphaSeq validation and 5 candidates for BLI validation, for each approach.

Results are shown in Figure 7: when mutating only the CDR we observed 60% successful candidates (i.e., an AlphaSeq reported affinity ≤ 1 uM), and in BLI we observed expression and binding from 4 of 5 top candidates, with 3 candidates within 10x of parental Trastuzumab affinity and 1 candidate with improved affinity over parental. When we optimized the entire CDRH1-CDRH3 window, we observed 23% success in AlphaSeq and 4/5 candidates with expression and binding in BLI, of which all 4 candidates were within 10x of parental affinity.

**Figure 7.**
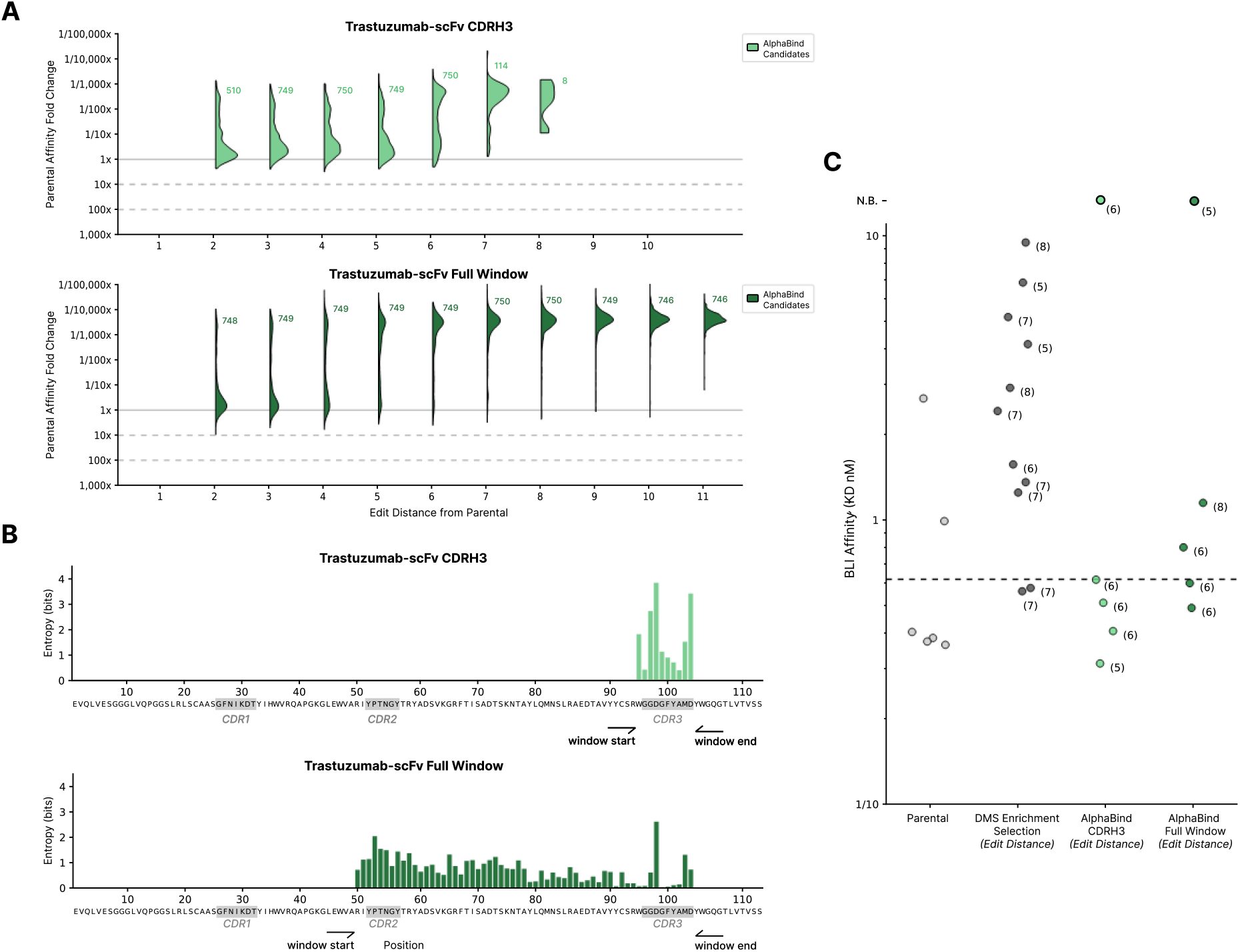
Optimization of Trastuzumab-scFv. **A:** AlphaBind models fine-tuned on mammalian display enrichment data for a combinatorial DMS (deep mutational scan) library of Trastuzumab-scFv CDRH3 variants, then used to generate candidates across two spans: only CDRH3 (concordant with the DMS data), and a larger mutable window covering CDRH2 through CDRH3. **B:** Sequence diversity as measured by Shannon entropy for the two spans is shown. Despite not being fine-tuned on any mutations in the framework regions, AlphaBind makes many edits in FR2 when allowed to mutate there. **C:** BLI validation results for parental Trastuzumab-scFv, DMS-optimized candidates selected by enrichment in the fine-tuning dataset, and AlphaBind-optimized candidates. 3 of 5 AlphaBind candidates with only mutations in CDRH3 were superior to the best directly observed in the fine-tuning dataset, and AlphaBind candidates in the full mutable window were comparable to those chosen by enrichment in the fine-tuning data despite being out of distribution with respect to sequences in the fine-tuning data. Edit distance for each candidate is shown in parentheses.

KinExA was used to confirm affinities for the top candidate from each optimization strategy, resulting in final relative affinity gains of 2.8x for the best CDRH3-only variant of Trastuzumab-scFv (from 7.43 pM to 2.74 pM, by KinExA) and 2.1x worse than parental for the best full-window variant of Trastuzumab-scFv (from 7.43 pM to 15.12 pM, by KinExA).

Overall, these results are comparable to those reported in the original study, which used *in silico* sequence proposal plus a purpose-built CNN classifier to create CDRH3 variants and found that 13/14 candidates bound HER2 (generally within 10x of parental) and 1/14 candidates had improved binding. Across both contexts the candidates optimized with AlphaBind showed good affinity with respect to the most-enriched candidates taken directly from the Mason et al. dataset used for fine-tuning: 3/5 (60%) of the AlphaBind candidates were better than any candidate from the training dataset when editing CDRH3. These data indicate that AlphaBind can optimize antibodies effectively with a different data format than what it was trained on, that AlphaBind-based optimization can substantially improve candidate quality over candidates assayed directly in the training dataset with minimal additional time and compute cost, and that our results are concordant with those previously demonstrated via an ML model built specifically for the Trastuzumab optimization task.

## Discussion

Across four separate parental systems—including humanized, humanoid and camelid antibodies— we have demonstrated that AlphaBind, with a combination of antibody-specific training data generated via unguided high-throughput affinity measurement, protein language models, and transfer learning from unrelated antibody-antigen affinity data, generates highly diverse antibody variants with substantial affinity improvements—although the scale of affinity improvement varied widely between 2x-74x for the four systems tested.

Exact comparisons between the scope of affinity gains from ML-guided optimizations are difficult due to widespread differences in parental antibodies, optimization methodology, and assessment techniques across experiments. However, we believe that these results are among the best reported for ML-guided optimization of antibody sequences, especially given only a single round of optimization rather than an iterative process. Angermueller, et al.^18^ recently reported a similar experiment to optimize VHH72, also using AlphaSeq training and validation datasets but utilizing an additional iteration of model training and optimization, and report 12/12 top candidates with improved affinity to SARS-CoV2-RBD by BLI, compared to 5/5 top candidates with improved affinity by BLI in the present study. Frey, et al.^15^ used an unguided method on Trastuzumab and reported 70% of designs expressed and bound target, compared to 80% (4/5) of our BLI tested designs that expressed and bound target. Li, et al.^19^ published a similar experiment optimizing naïve human Fabs targeting a SARS-CoV2 peptide, discovered by phage panning (i.e., similar to our discovery and optimization of AAB-PP489 but against a different target): they reported greater than 90% success rate of ML designs in AlphaSeq validation, including binders with AlphaSeq affinities up to 29x improved over any sequences in the training dataset. Finally, Krause, et al.^31^ recently utilized antibody-specific affinity data and language model guided design to optimize 5 parental antibodies binding CD40L and validated designs with SPR. They reported strong binders up to 8 mutations away from parental sequences, candidates with improved affinity for 4/5 parental antibodies, and an overall rate of approximately 40% of designed candidates with improved affinity compared to parental. In this study, we demonstrated binding on candidates with up to 11 mutations, and 10/10 (100%) tested candidates showed improved affinity by BLI.

As is evident in the comparisons above, the differences we observed in the scope of affinity gains across parental antibodies roughly accord with differences observed in other ML-guided antibody optimization papers, which report results ranging from a majority of sequences showing expression and some binding, but with affinity gains exceedingly rare^15,20^, to 90%+ of candidates showing improved binding compared to a parental antibody^19^. We speculate that these differences are primarily a function of how close each parental antibody is to a local optimum. It is notable that our work on AAB-PP489—an antibody that was optimized directly after discovery from a phage display library without *in vitro* or *in vivo* affinity maturation to drive it to a local optimum—had the best demonstrated improvement in affinity, at approximately 74x. Additionally, while our study demonstrates the effectiveness of AlphaBind across four distinct antibody-antigen systems, future work will be required to validate the model’s generalizability across a broader range of antibody classes, including complex cases such as antibodies targeting GPCRs, antibodies requiring conditional binding (e.g. pH dependence), antibodies which require broad cross-reactivity across homologs, paralogs, or strain variants of microbial targets, etc. Such studies will help further confirm AlphaBind’s utility in optimizing diverse antibody therapeutics.

Taken together, AlphaBind has demonstrated similar or improved performance compared to other recent studies across four different antibody optimization campaigns with a simple architecture, low training and optimization costs, significant transfer learning from unrelated antibody-antigen binding data, and a standardized codebase for easy adaptation to new contexts. While the affinity gains using this method varied widely between parental antibodies—from Pembrolizumab-scFv where almost all optimized candidates had improved affinity over the parental molecule, to Trastuzumab where only a small proportion of candidates had improved affinity—AlphaBind was able to generate candidates with improved affinity in each case. However, while we have demonstrated substantial affinity improvements across several parental antibodies, the most striking feature of these results is the breadth of primary sequence available among optimized candidates; for each of the parental antibodies studied, we identified thousands of variants with high affinity but highly diverse primary sequence. We anticipate that the ability to maintain or improve parental binding while opening a massive space of primary sequence will be critical to make best use of emerging capabilities in the prediction and engineering of non-affinity phenotypes including solubility, thermostability, and expression in mammalian cell culture systems—and we demonstrated this capability on an AlphaBind-derived variant of AAB-PP489, using the fine-tuned AlphaBind model to guide sequence-based improvements to predicted developability while further improving affinity. AlphaBind thus guided the optimization to a candidate with good expression, no observed aggregation, good thermostability, full ablation of sequence liabilities in the heavy chain, and femtomolar binding affinity by KinExA.

Finally, a key factor contributing to AlphaBind’s improved performance is the availability of large-scale (∼7.5 million AlphaSeq data points), high-quality, quantitative antibody-antigen interaction data for pre-training—allowing the model to capture general sequence-function relationships that enable transfer learning across parental antibodies. While this pre-training enhances AlphaBind’s ability to fine-tune effectively on specific datasets, it does not yet fully generalize across diverse binding landscapes. Achieving a truly generalizable model—capable of predicting optimal antibody sequences without the need for local fine-tuning—will require significantly larger and more diverse datasets, spanning a wider range of structural antibody-antigen systems, along with further pre-training. In contrast to structural models whose training datasets are growing very slowly as new solved structures accumulate, our corpus of quantitative molecular affinity data via AlphaSeq is large and growing rapidly, making the prospect of massive scaling of pretraining data entirely feasible in the near future. Looking ahead, such advancements in data collection and machine learning models will be critical for enabling zero-shot antibody engineering, where therapeutic antibodies can be designed entirely *in silico*. This will dramatically reduce the reliance on wet lab screening, accelerating early-stage drug discovery and making biologics development faster, more cost-effective, and more accessible.

## Materials and Methods

### AlphaSeq Data Generation

AlphaSeq analysis to generate large-scale affinity datasets for fine-tuning and large-scale candidate validation was performed as previously described^21,22^. Briefly, plasmids encoding yeast surface display cassettes were constructed and linearized for integration into the yeast genome. A 300bp oligonucleotide pool ordered from Twist Bioscience (South San Francisco, CA) was PCR amplified and inserted into each antibody backbone using Gibson assembly and subsequently PCR amplified. For the yeast library transformation, MATa and MATalpha AlphaSeq yeast were grown in YPAD media. Yeast cells were washed, resuspended, and incubated with the DNA library and unique barcodes using electroporation to pair each library member with multiple barcodes. DNA from the yeast libraries was extracted, and fragments containing the genes and associated DNA barcodes were PCR amplified. The amplified fragments were sequenced using a GridION sequencer from Oxford Nanopore Technologies (Oxford, UK) to link the DNA barcodes and antibody sequences. Library-on-library AlphaSeq assays were performed for each campaign by combining MATa and MATalpha libraries in YPAD media with a low concentration of Tween20 and incubating for 16 hours. Control yeast strains with known interaction affinities were included as standard controls. DNA fragments were amplified by qPCR with Illumina sequencing adaptors, which were then sequenced using Illumina NextSeq 500 (San Diego, CA). Sequencing data were analyzed to identify MATa and MATalpha barcode pairs, normalized based on haploid frequencies, and assigned estimated affinities using a linear regression derived from the control strains.

### AlphaBind Model Training

Briefly, AlphaBind takes as input ESM-2nv embeddings^14,32^ of both an antibody and a target sequence, then applies a transformer model with 4 attention heads and 7 layers trained to predict affinity with MSE loss (see Figure 2). AlphaBind network weights were pretrained using approximately 7.5 million measurements of AlphaSeq affinity data from unrelated antibody-antigen systems; due to a large quantity of relevant AlphaSeq data comprising antibodies targeting human TIGIT and SARS-CoV2-RBD, the full pretrained AlphaBind model was fine-tuned only for Pembrolizumab-scFv; a reduced AlphaBind model pretrained on approximately 1 million rows of unrelated antibody-antigen data was fine-tuned for VHH72 and AAB-PP489, in order to prevent information leakage.

After pretraining, antibody-specific AlphaBind models were fine-tuned using the AlphaSeq fine-tuning datasets for each parental antibody system. Fine-tuning each model took approximately 1 hour per parental antibody on a single H100 GPU, using a p5.48xlarge instance from Amazon Web Services (Seattle, WA). Models were trained for 100 epochs and the checkpoint with lowest validation score (using a uniform random 90/10 train/validation split) was chosen as the final model. We observed a relatively flat training curve after 40 epochs, suggesting that in routine use fewer epochs may be acceptable.

### AlphaBind Sequence Optimization and Candidate Selection

Each fine-tuned regression model was used to generate candidates for *in vitro* validation by stochastic greedy optimization of predicted binding affinity. Briefly, 60,000 trajectories for each parental antibody were optimized for 100 generations, during which a proposed sequence for each trajectory was randomly generated by masking 1-3 positions (not necessarily contiguous) within the mutational window (with the probability distribution: [1: 0.5, 2: 0.3, 3: 0.2]) in that trajectory’s current proposal and using ESM-2nv logits to sample mutations; if the new sequence had better predicted affinity according to the appropriate fine-tuned regression model, it was kept.

ESM-2nv embeddings and AlphaBind inference calls during this optimization process represent the main computational bottleneck for this work: optimization for each parental antibody was performed on AWS p5.48xlarge instances with 8 H100 GPUs and took approximately 5 hours per candidate optimization batch, for an overall cost of roughly $200 USD per parental antibody at the time of publication.

Once all trajectories were proposed and scored, we stratified the candidate sequences from all generations of all trajectories by their edit distance from the parental sequence, for the closed interval [2, 11], yielding 10 bins in total. From each bin, we selected the top 1,500 sequences by predicted affinity and screened them for predicted developability properties using TAP^5,6^, discarding sequences which had amber or red flags for any of the five TAP metrics. Finally, from each bin we selected up to 750 of the top remaining sequences by predicted affinity, yielding a total of approximately 7,500 selected sequences per target.

### Ablated AlphaBind Model Training and Optimization

Ablated AlphaBind models were constructed as follows: for ‘esm_warm’, training and optimization were identical to the full AlphaBind model, except that during the sequence proposal step of optimization, mutations were sampled at random rather than by using ESM-2nv logits, which necessitated 300 generations of optimization to accumulate sufficient candidates for validation. ‘esm_cold’ is similar to ‘esm_warm’, with the additional change that AlphaBind network weights were not pre-trained on unrelated antibody-antigen AlphaSeq data, and was likewise optimized for 300 generations; ‘ohe_warm’ and ‘ohe_cold’ were trained using a network architecture analogous to AlphaBind but using one-hot encoding rather than ESM-2nv for sequence embedding, and additionally comprised an ensemble of 5 models differentiated by random seed, due to very poor performance of single models; ‘ohe_warm’ was pre-trained on the same unrelated AlphaSeq data as the full AlphaBind model, and ‘ohe_cold’ was not pre-trained. ‘ohe_warm’ and ‘ohe_cold’ both generated sufficient candidates for validation within 100 generations due to high acceptance rates. We provide the AlphaBind GitHub repository (https://github.com/A-Alpha-Bio/alphabind) for additional technical details on ablated model training and optimization.

### Biolayer Interferometry (BLI) and Kinetic Exclusion Assays (KinExA)

Sequences for BLI validation were chosen by a similar method as above, incorporating an additional filtering criterion of zero introduced liabilities compared to the parental sequence, scored using software from NaturalAntibody (Szczecin, Poland)^28^. The top 5 sequences by predicted affinity were then selected for BLI validation. In addition to our experimental candidates, 3 replicates of the parental antibody sequence were submitted for expression and BLI at Twist Bioscience. All 3 parental replicates for Pembrolizumab-scFv had insufficient expression to enable BLI measurements.

In order to confirm measured affinity values and establish better quantitative comparisons of parental and variant affinity for the best AlphaBind candidates, single-point BLI at Twist was followed by either multi-point BLI (for VHH72 variants), or Kinetic Exclusion Assay (KinExA) ^33^ where single-point BLI indicated an off-rate near or below the limit of detection (i.e., for the best AAB-PP489 and Trastuzumab-scFv variants from our initial optimization campaigns, as well as AAB-PP3115 and AAB-PP3117). See the Supplemental Materials for additional information on BLI and KinExA methods and results.

## Supporting information

Supplemental Methods & Results

## Code and Data Availability

Code used to train fine-tuned AlphaBind models and to conduct sampling, AlphaSeq fine-tuning and validation datasets, notebooks to reproduce the results and figures in this study, and trained AlphaBind model weights are all included in an open-source GitHub repository under the MIT license [cite link].

## Acknowledgements

We would like to acknowledge Dasha Krayushkina, Leah Homad, Jacob Brockerman, and the rest of the Technical Operations team from A-Alpha Bio for their assistance with the AlphaSeq and BLI data generation for this work. We would also like to acknowledge the generous additional support provided by the following Nvidia collaborators who assisted with technical details and application of BioNeMo ESM-2nv for AlphaBind: Darren Hsu, John Judge, Thomas Grilli, Vega Shah, and Neha Tadimeti.

