## Supplemental Methods & Results for "AlphaBind, a Domain-Specific Model to Predict and Optimize Antibody-Antigen Binding Affinity"

**Processing and Transformation of Trastuzumab DMS Data**

For training data derived from deep mutational scanning (DMS), we processed raw data from Mason et al. containing CDRH3 variants of Trastuzumab selected under antigen-positive and antigen-negative conditions using a mammalian display assay^1^. After filtering for sequence quality and removing duplicates, we calculated enrichment ratios by comparing adjusted fractions between conditions, using pseudocounts to account for sequences observed in only one condition. The CDRH3 sequences were then embedded within the complete Trastuzumab-scFv framework (consisting of VH and VL domains connected by a (G4S)3 linker) to generate full-length sequences. To convert enrichment ratios into a format compatible with our training pipeline, we performed z-score normalization of log-enrichment ratios and shifted the distribution to approximate binding affinity measurements, similar to the distribution of binding affinities in our experimental dataset—specifically, we flipped the sign of z-scores so that lower numbers indicate stronger binding, as in affinity values, and re-centered the distribution at 2 (i.e., 100nM) to approximate common affinity ranges seen in the AlphaSeq pretraining data.

**Affinity-Guided** **Developability Engineering**

Starting from AAB-PP3115, V and J germline calls and CDR regions were annotated with ANARCI according to Chothia nomenclature^2^, and potential sequence liabilities assessed with NaturalAntibody^3^. Two liability motifs in CDRH3 were identified: a DS isomerization motif at Chothia positions 100-100A and a DP fragmentation motif at Chothia positions 101-102. The trained AlphaBind regressor from our AAB-PP489 optimization campaign was used to predict binding affinities against human TIGIT for all combinations of mutations at those sites (19^4^ = 130,321 candidates). Candidates with predicted affinities within 0.5 logs of AAB-PP3115 and no liabilities were retained. Nine positions within the previously defined mutable window (see Table S1) were also identified as non-germline residues; for each retained candidate from the previous step, we further generated all 512 (2^9^) germline-reverted candidates. Again, candidates were retained if the fine-tuned AlphaBind regressor predicted an affinity within 0.5 logs of AAB-PP3115. A subset of 11 candidates with no sequence liabilities, predicted immunogenicity < 10 as annotated by NaturalAntibody, and all green flags from TAP^4^ were chosen for BLI validation based on predicted affinity, extent of germline reversion, and sequence diversity. The parental candidate AAB-PP3115, as well as the best candidate with all nine positions reverted to germline and no liabilities, were also included as comparators for a total of 13 sequences.

**Single-Point Biolayer Interferometry (BLI)**

Protein expression, purification and measurements for single-point BLI for AlphaBind candidates from the three main optimization campaigns plus Trastuzumab-scFv were performed by Twist Bioscience (South San Francisco, CA). Single-point BLI was performed using an Octet RH96 instrument from Sartorius (Göttingen, Germany). Results are shown in Table S3 and S4 below.

**BLI & Protein Analytics for Affinity-Guided Developability Engineering**

Single-point BLI for AAB-PP3115 and its variants was performed at A-Alpha Bio using the Gator Prime instrument (I&L Biosystems, Troisdorf, Germany). Anti-Human IgG Fc probes were loaded with target proteins at 10μg/mL final concentration in BLI sample buffer (1x PBS, 0.4%BSA, 0.02% PS20) for 180 seconds, washed with buffer in baseline wells for 30 secs, associated with human TIGIT protein (TIT-H52H5 from ACROBiosystems, Newark, DE) at 50nM concentration for 120 seconds for K_on_ measurements, then moved back to baseline well for 120 second dissociation to determine K_off_ measurements. The binding affinity (K_D_) was determined with the Gator analysis software.

BLI was also used to determine the expression titer using the Gator Prime instrument in a quantitation assay format. Probes were loaded with 200 uL of clarified ExpiCHO supernatant, and on-rates were compared to a standard curve of a similarly formatted control scFv-Fc protein to determine expression titer using Gator analysis software.

Protein purity was determined using an Agilent 1260 Infinity II HPLC (Agilent Technologies, Santa Clara, CA). Up to 25 ug purified protein was run over a tandem Advance Bio SEC 200A 1.9um 4.6x30mm Guard and Advance Bio SEC 200A 1.9um 4.6x150mm column set at 0.35ml/min in 50mM KH_2_PO_4_/150mM K_2_HPO_4_ running buffer. Agilent ChemStation software was used to acquire A280 values and determine retention time and area integration, and results are reported as % Main Peak.

Differential Scanning Fluorimetry (DSF) was performed on a QuantStudio 3 Real-Time PCR instrument (Applied Biosystems, Thermo Fisher Scientific Inc., Waltham, MA). Briefly, 5ug of purified protein samples were prepared in PBS using Protein Thermal Shift Dye Kit. Melting temperature (T_m_) was determined using Protein Thermal Shift Software v1.4 (Applied Biosystems) using the mean derivative of the earliest protein domain folding event of samples run in quadruplicate.

Results for all assays are reported in Table S5.

**Multi-Point BLI**

Multi-point BLI was used to confirm the binding affinity (K_D_) for VHH72 parental and its best-performing variant from single-point BLI, using a Gator Prime instrument (I&L Biosystems, Trosdorf, Germany). Anti-Human IgG Fc probes were loaded with target proteins at 10μg/mL final concentration in BLI sample buffer (1x PBS, 0.4%BSA, 0.02% PS20) until response on all probes met threshold of 1nm shift, washed with buffer in baseline wells for 30 secs, associated with titrated concentrations of SARS-CoV-2 S protein RBD (SPD-C52H3 from ACROBiosystems, Newark, DE) for 5 min for K_on_ measurements, then moved back to baseline well for 5-10 min dissociation. Affinity was determined with the Gator analysis software.

Multi-point BLI results are shown in Table S6 below.

**Kinetic Exclusion Assay (KinExA)**

K_D_ measurements were performed at room temperature using a KinExA 4000 (Sapidyne Instruments Inc., Boise ID) as previously described^5^. Briefly, binding curves were generated by measuring the concentration of free antibody (scFv-FC or IgG) in a series of samples containing a constant concentration of antibody with a 2-fold titration series of the antigen and allowed to come to equilibrium at room temperature (minimum of 12 hours). Two binding curves were run at different constant concentrations of antibody and run on the KinExA instrument. Analysis was performed using data from both equilibrium curves with Sapidyne Instruments’ KinExA Pro software (version 4.7.4) n-curve module.

We set up the KinExA 4000 to measure free antibody in equilibrated samples by capturing unbound antibody on a solid phase coated with antigen and detecting the amount of antibody on the solid phase with a fluorescent secondary probe. The observation column was loaded with solid phase and equilibrated antibody-antigen mixture was flowed over the solid phase at a rate of 0.25 mL/min. The volume varied (0.05 mL to 8.0 mL) according to specific antibody concentration to achieve signals in the optimal working range. Post sample loading, 500 uL of 500 ng/mL secondary corresponding to the format of the antibody (either anti-IgG H+L or IgG FCg) was flowed over the column at 0.25 mL/min to label the captured free antibody and signals were read once per second (Alexa Fluor® 647 AffiniPure Goat Anti-Human IgG FCγ (Part # 109-605-008); Alexa Fluor® 647 AffiniPure Goat Anti-Human IgG H+L (Part # 109-605-003 ) Jackson ImmunoResearch Inc., West Grove, PA) An antibody only control was used to determine the maximum signal for each equilibration curve. Analysis on the KinExA Pro software provides 95% confidence intervals in addition to the best-fit KD value which are reported in the results.

The solid phase, azlactone activated beads, (Sapidyne Part#444110) was coated with the target antigen, HER2 or TIGIT (HE2-H5225, TIT-H52H5 ACROBiosystems, Newark, DE) at a concentration of 20ug/mL following instructions (HG209). Briefly, antigen was diluted down to 20ug/mL in carbonate buffer (Sapidyne Part# 2S6320) and added to the azlactone activated beads. Beads were incubated on a rotator at room temperature for 1 hour before removing the antigen solution and replacing with 1mL Tris buffer + BSA (Sapidyne Part# 2T5500) to block the beads for an additional hour. Beads were stored at 4C until use.

KinExA results are shown in Table S7 below.

**Supplemental Results**

**Table S1 (separate Microsoft Excel file): Summary of Antibody Optimization Campaigns**

**Table S2: Summary of AlphaSeq Datasets**

| **Dataset** | **# Rows** | **Description** | **Variants** |
| --- | --- | --- | --- |
| aab-pp489_fine-tuning_alphaseq.csv | 26,913 | Fine-tuning dataset for AAB-PP489 | Unguided variants near parental |
| pembrolizumab-scfv_fine-tuning_alphaseq.csv | 29,897 | Fine-tuning dataset for Pembrolizumab-scFv | Unguided variants near parental |
| vhh72_fine-tuning_alphaseq.csv | 29,988 | Fine-tuning dataset for VHH72 | Unguided variants near parental |
| aab-pp489_validation_alphaseq.csv | 37,346 | Validation dataset for AAB-PP489 | Guided variants, AlphaBind + ablated models |
| pembrolizumab-scfv_validation_alphaseq.csv | 37,489 | Validation dataset for Pembrolizumab-scFv | Guided variants, AlphaBind + ablated models |
| vhh72_validation_alphaseq.csv | 37,070 | Validation dataset for VHH72 | Guided variants, AlphaBind + ablated models |
| trastuzumab-scfv-cdr_validation_alphaseq.csv | 3,681 | Validation dataset for Trastuzumab-scFv (CDRH3) | Guided variants, AlphaBind |
| trastuzumab-scfv-full_validation_alphaseq.csv | 7,540 | Validation dataset for Trastuzumab-scFv (full) | Guided variants, AlphaBind |

**Table S3: Single-Point BLI Results, Main AlphaBind Campaigns**

| Ab ID | Ab Capture (nm) | Binding Response (nm) | K_D_ (M) | K_on_(1/Ms) | K_off_(1/s) | R^2^ |
| --- | --- | --- | --- | --- | --- | --- |
| aab-pp489_wt_0 | 1.08 | 0.29 | **1.54E-09** | 4.78E+05 | 7.38E-04 | 0.96 |
| aab-pp489_wt_1 | 1.04 | 0.29 | **9.80E-10** | 5.35E+05 | 5.24E-04 | 0.97 |
| aab-pp489_wt_2 | 1.09 | 0.31 | **1.25E-09** | 4.67E+05 | 5.82E-04 | 0.97 |
| aab-pp489_alphabind_1500_d4 | 0.74 | 0.22 | **2.11E-10** | 5.65E+05 | 1.19E-04 | 0.95 |
| aab-pp489_alphabind_1502_d4 | 1.08 | 0.32 | **3.89E-10** | 4.32E+05 | 1.68E-04 | 0.97 |
| aab-pp489_alphabind_2254_d5 | 1.1 | 0.32 | **5.16E-10** | 4.08E+05 | 2.10E-04 | 0.97 |
| aab-pp489_alphabind_3750_d7 | 0.95 | 0.28 | **2.16E-10** | 4.62E+05 | <1.00E-04 | 0.98 |
| aab-pp489_alphabind_5252_d9 | 1.04 | 0.3 | **2.33E-10** | 4.30E+05 | <1.00E-04 | 0.98 |
| vhh72_wt_0 | 0.91 | 0.51 | **1.40E-08** | 7.77E+05 | 1.09E-02 | 0.95 |
| vhh72_wt_1 | 0.9 | 0.51 | **1.67E-08** | 6.60E+05 | 1.10E-02 | 0.95 |
| vhh72_wt_2 | 0.94 | 0.54 | **1.62E-08** | 6.51E+05 | 1.06E-02 | 0.95 |
| vhh72_alphabind_3000_d6 | 1.02 | 0.57 | **1.26E-08** | 9.07E+05 | 1.14E-02 | 0.93 |
| vhh72_alphabind_5284_d9 | 0.99 | 0.61 | **2.73E-09** | 5.19E+05 | 1.42E-03 | 0.97 |
| vhh72_alphabind_6056_d10 | 0.93 | 0.58 | **1.38E-09** | 6.92E+05 | 9.58E-04 | 0.98 |
| vhh72_alphabind_6786_d11 | 1.24 | 0.72 | **6.22E-09** | 4.63E+05 | 2.88E-03 | 0.98 |
| vhh72_alphabind_6808_d11 | 0.95 | 0.53 | **2.46E-09** | 7.34E+05 | 1.81E-03 | 0.98 |
| pembrolizumab-scfv_alphabind_1171_d3 | 0.2 | 0.02 | **1.67E-08** | 2.86E+04 | 4.79E-04 | 0.92 |
| pembrolizumab-scfv_alphabind_3153_d6 | 1.23 | 0.19 | **3.54E-08** | 2.26E+05 | 7.99E-03 | 0.96 |
| pembrolizumab-scfv_alphabind_4784_d8 | 1.19 | 0.31 | **1.18E-08** | 8.45E+05 | 9.95E-03 | 0.97 |
| pembrolizumab-scfv_alphabind_6160_d10 | 1.12 | 0.21 | **2.30E-08** | 3.56E+05 | 8.18E-03 | 0.94 |
| pembrolizumab-scfv_alphabind_6203_d10 | 1.21 | 0.11 | **5.45E-08** | 1.08E+05 | 5.86E-03 | 0.94 |

**Table S4: Single-Point BLI Results, Trastuzumab-scFv**

| Ab ID | Ab Capture (nm) | Binding Response (nm) | K_D_ (M) | K_on_(1/Ms) | K_off_(1/s) | R^2^ |
| --- | --- | --- | --- | --- | --- | --- |
| trastuzumab-scfv-cdr_wt_0 | 1.17 | 0.8 | **3.64E-10** | 2.75E+05 | < 1.00E-04 | 0.96 |
| trastuzumab-scfv-cdr_wt_1 | 1.18 | 0.8 | **3.85E-10** | 2.60E+05 | < 1.00E-04 | 0.97 |
| trastuzumab-scfv-cdr_wt_2 | 1.12 | 0.51 | **9.92E-10** | 1.01E+05 | < 1.00E-04 | 1 |
| trastuzumab-scfv-full_wt_0 | 1.19 | 0.68 | **2.68E-09** | 2.77E+05 | 7.42E-04 | 0.97 |
| trastuzumab-scfv-full_wt_1 | 1.09 | 0.78 | **3.74E-10** | 2.68E+05 | < 1.00E-04 | 0.97 |
| trastuzumab-scfv-full_wt_2 | 1.09 | 0.75 | **4.04E-10** | 2.48E+05 | < 1.00E-04 | 0.97 |
| dms_candidate_3994_d6 | 1.33 | 0.8 | **1.57E-09** | 2.66E+05 | 4.18E-04 | 0.97 |
| dms_candidate_4178_d5 | 1.19 | 0.36 | **4.16E-09** | 2.07E+05 | 8.62E-04 | 0.97 |
| dms_candidate_5272_d7 | 1.22 | 0.24 | **1.25E-09** | 8.00E+04 | < 1.00E-04 | 0.99 |
| dms_candidate_7234_d5 | 1.26 | 0.46 | **6.85E-09** | 1.17E+05 | 8.03E-04 | 1 |
| dms_candidate_9082_d8 | 1.25 | 0.79 | **2.92E-09** | 2.98E+05 | 8.72E-04 | 0.97 |
| dms_candidate_9091_d7 | 1.24 | 0.41 | **1.36E-09** | 2.19E+05 | 2.97E-04 | 0.96 |
| dms_candidate_9208_d7 | 1.48 | 0.46 | **5.18E-09** | 1.05E+05 | 5.41E-04 | 0.99 |
| dms_candidate_9285_d8 | 1.39 | 0.17 | **9.48E-09** | 4.60E+04 | 4.36E-04 | 9.9 |
| dms_candidate_9294_d7 | 1.31 | 0.76 | **5.77E-10** | 1.73E+05 | < 1.00E-04 | 0.99 |
| dms_candidate_9765_d7 | 1.43 | 0.07 | **2.42E-09** | 4.14E+04 | < 1.00E-04 | 0.97 |
| dms_candidate_10075_d7 | 1.25 | 0.73 | **5.62E-10** | 3.16E+05 | 1.77E-04 | 0.95 |
| trastuzumab-scfv-cdr_alphabind_2019_d5 | 1.33 | 0.89 | **3.13E-10** | 3.20E+05 | < 1.00E-04 | 0.95 |
| trastuzumab-scfv-cdr_alphabind_2764_d6 | 1.33 | 0.76 | **6.17E-10** | 1.62E+05 | < 1.00E-04 | 0.99 |
| trastuzumab-scfv-cdr_alphabind_2773_d6 | 1.24 | 0.77 | **5.12E-10** | 1.95E+05 | < 1.00E-04 | 0.98 |
| trastuzumab-scfv-cdr_alphabind_2778_d6 | 1.3 | 0.84 | **4.07E-10** | 2.46E+05 | < 1.00E-04 | 0.97 |
| trastuzumab-scfv-full_alphabind_3244_d6 | 1.14 | 0.62 | **6.00E-10** | 1.67E+05 | < 1.00E-04 | 0.99 |
| trastuzumab-scfv-full_alphabind_3282_d6 | 1.22 | 0.5 | **8.02E-10** | 1.25E+05 | < 1.00E-04 | 0.99 |
| trastuzumab-scfv-full_alphabind_3313_d6 | 1.1 | 0.55 | **4.91E-10** | 2.04E+05 | < 1.00E-04 | 0.98 |
| trastuzumab-scfv-full_alphabind_4628_d8 | 1.3 | 0.31 | **1.15E-09** | 8.68E+04 | < 1.00E-04 | 1 |

**Table S5 (separate Microsoft Excel file): BLI & Protein Analytics, AAB-PP3115 & Variants**

**Table S6: Multi-Point BLI Results**

| **Ab ID** | **Target** | **K_D_(M)** | **K_on_(1/Ms)** | **K_off_(1/s)** | **Full R^2^** | **K_on_ Error** | **K_off_ Error** |
| --- | --- | --- | --- | --- | --- | --- | --- |
| **VHH72** | RBD | **1.68E-08** | 8.03E+05 | 1.35E-02 | 0.968 | 7.22E+03 | 3.61E-05 |
| **vhh72_alphabind_6056_d10** | RBD | **1.17E-09** | 6.10E+05 | 7.12E-04 | 0.992 | 1.34E+03 | 1.25E-06 |

**Table S7: KinExA Results**

| **Ab ID** | **Format** | **Target** | **K_D_** | **95% CI** | **%error** |
| --- | --- | --- | --- | --- | --- |
| **AAB-PP489** | ScFv-FC | TIGIT | **56.51 pM** | 44.32pM-70.83pM | 3.20 |
| **aab-pp489_alphabind_1500_d4** | ScFv-FC | TIGIT | **766.25 fM** | 250.64fM-1.57pM | 3.28 |
| **AAB-PP3115** | IgG | TIGIT | **1.04pM** | 571fM-1.67pM | 2.23 |
| **AAB-PP3117** | IgG | TIGIT | **309.81 fM** | 111.57-579.03 fM | 1.92 |
| **Trastuzumab-scFv** | ScFv-FC | HER2 | **7.34pM** | 5.89pM-9.07pM | 1.88 |
| **trastuzumab-scfv-full_alphabind_3313_d6** | ScFv-FC | HER2 | **15.12pM** | 11.8pM-19.11pM | 1.5 |
| **trastuzumab-scfv-cdr_alphabind_2019_d5** | ScFv-FC | HER2 | **2.77 pM** | 1.64pM-4.52pM | 4.28 |
